# Microbial exometabolite responses to root exudates from tallgrass prairie plants

**DOI:** 10.64898/2026.05.29.728892

**Authors:** Vlastimil Novak, Markus de Raad, Ananya Vittaladevuni, Vadant Saraf, Meghan Midgley, Chelsea Hernandez, Nicholas A. Barber, Trent R. Northen

## Abstract

Root exudates influence soil microbial community assembly and function, yet plant species-specific effects on microbially-driven metabolic transformations remain insufficiently characterized under controlled conditions. Here, we present a multi-omic dataset of exudate–microbe interactions across six plant species, including tallgrass prairie species (*Helianthus pauciflorus, Carex brevior, Eragrostis* spectabilis, *Astragalus canadensis*, and *Panicum virgatum*) and a model grass (*Brachypodium distachyon*), generated using an *in vitro* incubation experiment. Sterile root exudates were used to amend a soil-mimicking medium inoculated with a native tall-grass prairie soil microbiome. The dataset includes full-length 16S rRNA gene sequencing for microbial community profiling, untargeted LC-MS/MS metabolomics for exometabolomic profiling, and optical density measurements of microbial growth. We describe the axenic plant growth protocols, experimental design, data acquisition, processing workflows, and technical validation. This dataset provides a resource for investigating microbially mediated exometabolite transformations across diverse plant species to help understand rhizosphere processes, microbiome assembly, and ecosystem function.

## 1. Background & Summary

Plant root exudates are a mixture of compounds released into the rhizosphere, where they serve as microbial substrates and influence soil community assembly^1,2^. Their composition varies across plant species^3,4^, genotypes^5^, and abiotic factors^6^, contributing to differences in microbial activity. In ecosystems such as tallgrass prairies, which combine species-rich plant and microbial communities with substantial belowground carbon stocks, exudates are an important driver of soil microbial processes^7,8^. However, direct links between specific exudates and microbial transformations of these exometabolites remain difficult to resolve due to soil complexity^9^. This limitation motivates the use of controlled experimental systems that enable defined exudate inputs and paired measurements of microbial composition and metabolic outputs^10^.

In this study (**Fig. 1**), we conducted controlled *in vitro* microbial incubations with root exudates from multiple understudied tallgrass prairie species often found in plant communities: *Helianthus pauciflorus* (showy sunflower, Asteraceae), *Carex brevior* (plains oval sedge, Cyperaceae), *Eragrostis spectabilis* (purple lovegrass, Poaceae), *Astragalus canadensis* (Canada milkvetch, Fabaceae), and *Panicum virgatum* (switchgrass, Poaceae), alongside the model grass *Brachypodium distachyon* (stiff brome, Poaceae), serving as a non-native comparison with prairie plants. Plant axenic growth protocols were developed to enable sterile root exudate collection. To support microbial growth and exudate utilization, we formulated an assay medium based on the Northen Lab Defined Medium (NLDM), a mixture of common soil metabolites used to assess bacterial substrate utilization^11,12^. Here, NLDM was applied at reduced organic carbon concentration to approximate soil metabolite background levels^13^, with root exudates added as additional substrates^1^. The different exudate assay media were inoculated with a natural microbial community from prairie soil or maintained as sterile controls.

**Fig. 1:**
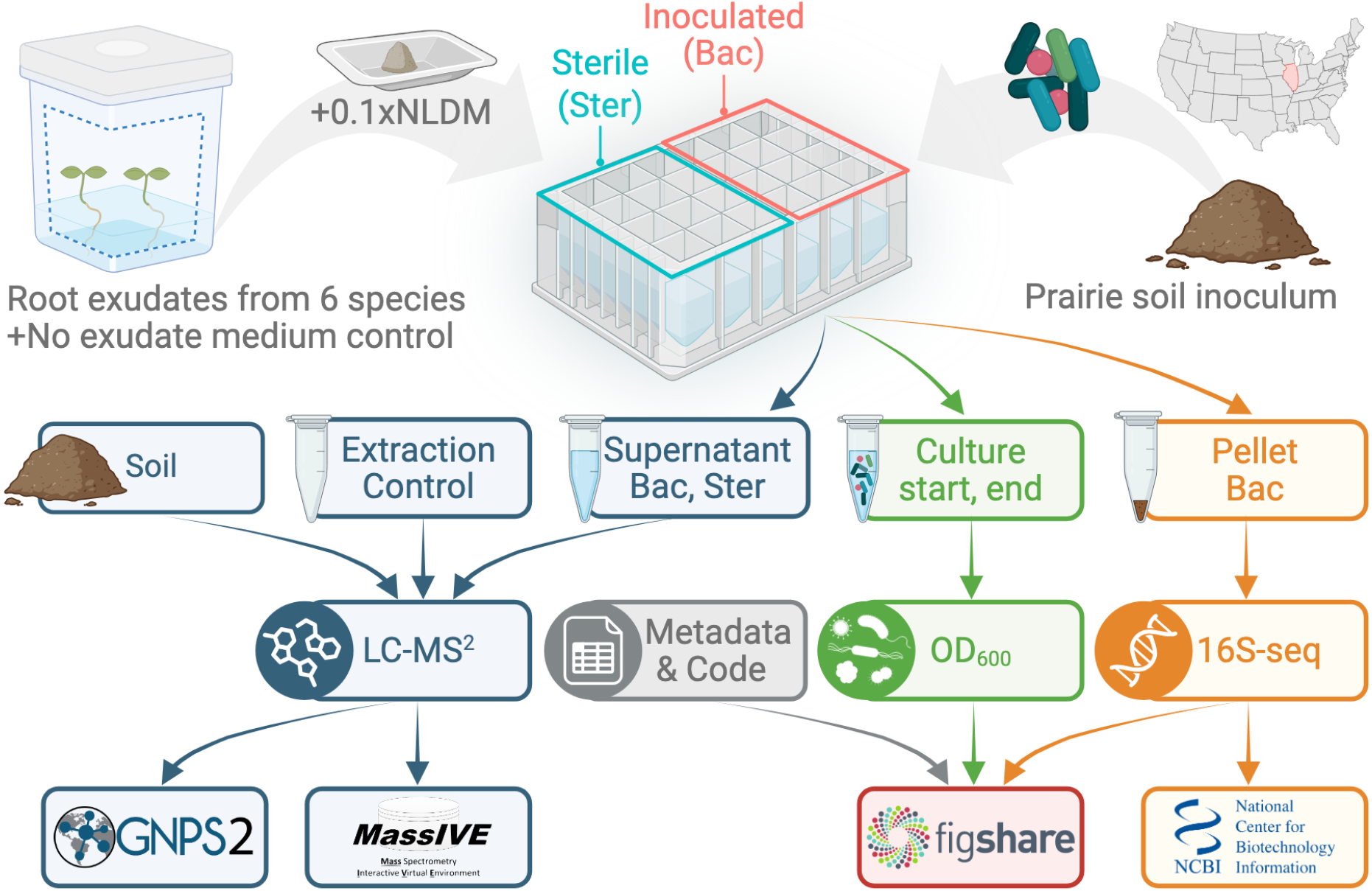
Experimental overview of exudate-microbiome experiment. The root exudates were collected from six plants: *Helianthus pauciflorus, Carex brevior, Eragrostis spectabilis, Astragalus canadensis, Panicum virgatum*, and *Brachypodium distachyon*. We also used no exudate medium as control. Additional labile carbon was supplied as 0.1x diluted NLDM to mimic the soil background. The resulting assay media was inoculated (Bac) with extract from prairie soil (Lisle, Illinois, USA) or left sterile (Ster). The supernatant, soil, and extraction control samples were analyzed by LC-MS/MS, the data was analyzed in GNPS2 and were archived in MassIVE and GNPS2. Inoculated culture pellets were measured by full-length 16S rRNA sequencing and archived on Figshare and NCBI. Culture OD_600_ was measured and deposited in Figshare, alongside metadata, code, and additional files.

The resulting dataset integrates three complementary data types (**Fig. 1**). First, untargeted LC-MS/MS metabolomics captures the chemical composition of prairie soil alongside culture supernatants, including both plant-derived metabolites and microbially transformed products following growth in different exudate environments. Second, full-length 16S rRNA gene sequencing provides high-resolution profiling of microbial community composition. Third, optical density measurements (OD_600_) provide an estimate of microbial growth across treatments. Together, these measurements enable investigation of relationships between plant-derived substrates, microbial community structure, and metabolic outcomes.

Public platforms such as Global Natural Products Social Molecular Networking (GNPS) ^14^ and National Center for Biotechnology Information (NCBI)^15^ facilitate the sharing and reuse of data. In accordance with this, all raw and processed data, along with metadata and analysis workflows, have been deposited in public repositories, including GNPS2/MassIVE for metabolomic data and NCBI and Figshare for sequencing data and associated files (**Fig. 1**). The dataset is accompanied by detailed descriptions of the experimental design, sample processing, and data acquisition to facilitate reuse in accordance with the FAIR (Findable, Accessible, Interoperable, and Reusable) principles^16^. Additionally, we have followed recently developed STREAMS reporting guidelines for environmental microbiome studies^17^.

This resource supports multiple avenues of secondary analysis. For example, it enables comparative studies of how plant species identity influences microbial community assembly under standardized conditions, as well as investigations of microbially mediated transformation of exometabolites. The paired metabolomic and microbiome data enable exploration of associations between specific taxa and metabolic features, development and benchmarking of integrative computational methods, and assessment of microbial responses to exometabolite inputs and their stability.

## 2. Methods

### 2.1. Plant exudate production

The six plant species included: *H. pauciflorus* (ncbitaxon:382521), *C. brevior* (ncbitaxon:314401), *E. spectabilis* (ncbitaxon:888271), *A. canadensis* (ncbitaxon:20408), *P. virgatum* (ncbitaxon:38727), and *B. distachyon* Bd21-3 (ncbitaxon:15368)^18^. We obtained seeds from Prairie Moon Nursery (Winona, MN, USA) with catalog IDs: *H. pauciflorus* #HEL48F, *C. brevior* #CAR39G, *E. spectabilis* #ERA02G, *A. canadensis* #AST52F, *P. virgatum* #PAN04G, while *B. distachyon* Bd21-3 seeds were from John Vogel (Joint Genome Institute, CA, USA).

Seed sterilization, stratification, and germination followed the *B. distachyon* protocol^10^, with a few modifications for certain species to achieve sterility (**Table 1**). Seeds of *B. distachyon* and *C. brevior* were de-husked, while others were not. *H. pauciflorus* and *P. virgatum* were treated with 3% H_2_O_2_ for 1h. All species were then sterilized with 70% ethanol for 30s, followed by 5 min in 6% sodium hypochlorite, and 5 washes with sterile Milli-Q water. Seeds were then stratified on 1.5% (w/v) Phytoagar (40100072, PlantMedia™) and ½ MS basal salts (M524, PhytoTech Lab) at 4 °C in the dark for species-optimized period of 3 days for *B. distachyon* and 14 days for other species. Germination was induced by placing plates vertically (~80° angle) in a growth chamber (CU36L4, Percival Scientific Inc., Perry, IA, USA). The chamber was set to 22 °C for the duration of the experiments, with a photosynthetic photon flux density of 150 μmol/m^2^/s and a 12-hour photoperiod. The germination period ranged from 3 to 21 days, depending on growth rate, to ensure sufficient size for transfer to magenta boxes (**Table 1**).

**Table 1:** Production of sterile root exudates.

| Group code | Plant name | Seed de-husking | Seed in 3% H<sub>2</sub>O<sub>2</sub> for 1h | Seeds in 70% EtOH for 30s + 6% NaOCl for 5min | Stratification time at 4°C (days) | Germination time (days) | Number of plants per box at t<sub>0</sub> | Growth time in boxes until harvest (days) | Number of plants per box at harvest |
| --- | --- | --- | --- | --- | --- | --- | --- | --- | --- |
| BRADIS | *Brachypodium distachyon* | TRUE | FALSE | TRUE | 3 | 3 | 3 | 21 | 2-3 |
| HELPAU | *Helianthus pauciflorus* | FALSE | TRUE | TRUE | 14 | 18 | 3 | 49 | 2-3 |
| CARBRE | *Carex brevior* | TRUE | FALSE | TRUE | 14 | 15 | 3 | 43 | 2-3 |
| ERASPE | *Eragrostis spectabilis* | FALSE | FALSE | TRUE | 14 | 12 | 3 | 38 | 2-3 |
| ASTCAN | *Astragalus canadensis* | FALSE | FALSE | TRUE | 14 | 10 | 4 | 55 | 2-4 |
| PANVIR | *Panicum virgatum* | FALSE | TRUE | TRUE | 14 | 21 | 1 | 47 | 1 |
| NoExu | No plant medium control | NA | NA | NA | NA | NA | 0 | NA | NA |

Footnote: NA - not applicable.

Germinated seedlings were transferred to culture boxes (#C2100, PhytoTech Lab) containing 55 mL of ½ MS basal salts (M524, PhytoTech Lab), covered with vented lids (#C2110, PhytoTech Lab), and placed in a growth chamber. To prevent overcrowding, different numbers of seedlings were transferred based on species (**Table 1**). Seedlings were thinned one week after transfer; to retain healthy plants, while poorly developed seedlings were removed. Growth durations (21–55 days) were tailored to when plants reached the top of the boxes. For exudate collection, we selected 3-5 boxes with the most comparable plant growth and pooled the spent media to obtain a representative exudate sample. Boxes with ½ MS salts without plants were incubated for 55 days in a chamber, and the medium was collected as a baseline. To verify sterility, 50 uL of pooled media were incubated on LB plates for 7 days at 30ºC in the dark. The remaining media was filtered through a 0.2 µM PES membrane and stored at 4ºC until use.

### 2.2. Preparation of assay media from root exudates

The assay media were prepared by adding components of NLDM (organics, salts, vitamins, and minerals)^11,12^ to the exudate solution or the no-exudate control (NoExu), with the NLDM organics concentration set to 0.1x (104.2 ppm C) (**Table 2**). The adjustment of NLDM organics concentration was driven by our objective to create a growth-supporting medium while maintaining metabolite concentrations similar to those in native soil^13^. The target pH of the media was 7.0 ± 0.1. For most plant species, the media did not need adjustment after ingredients mixing; only CARBRE and ERASPE were adjusted with 2 M NaOH. The assay media was filtered via a 0.2 µM PES membrane.

**Table 2:**
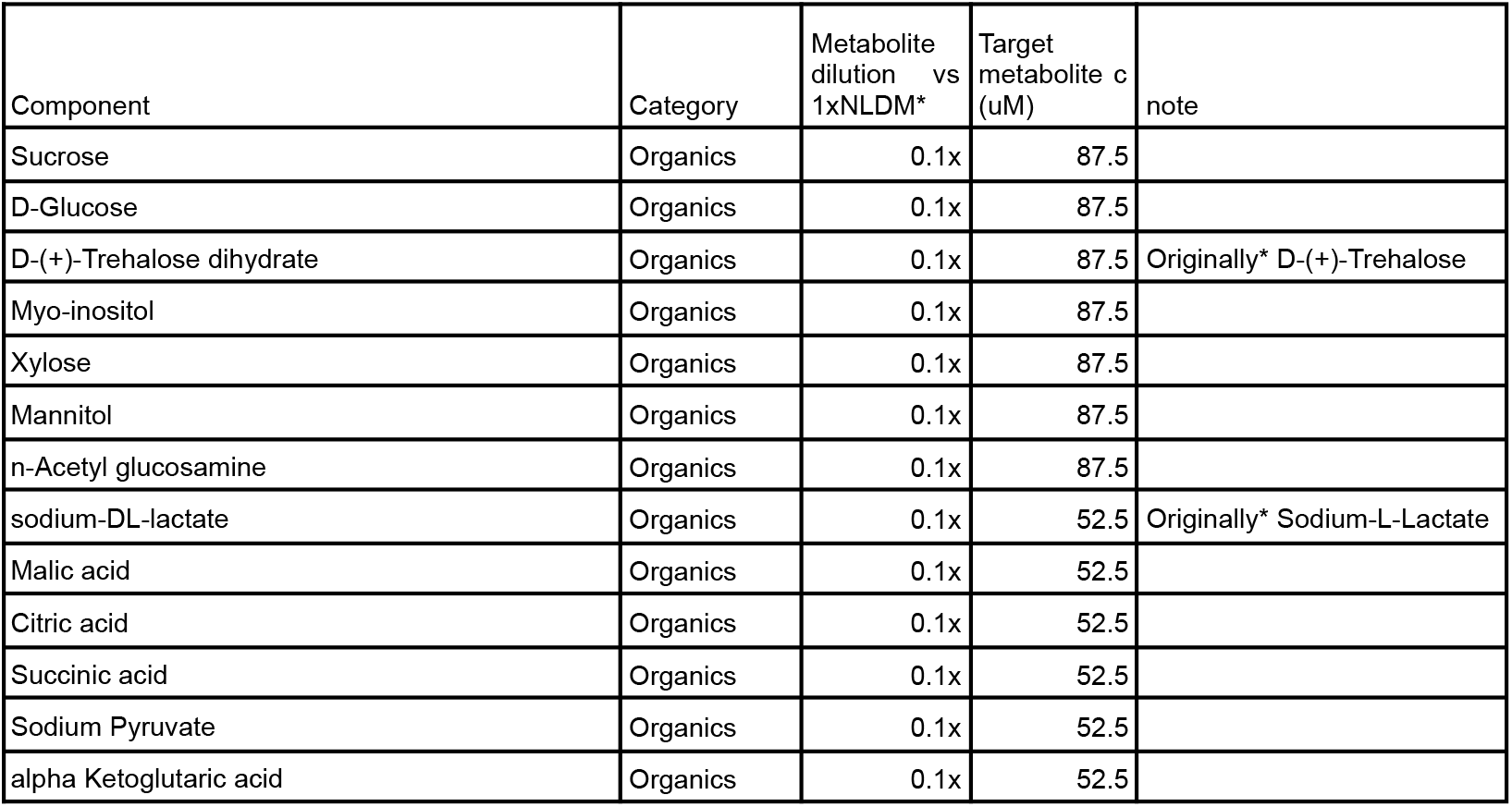
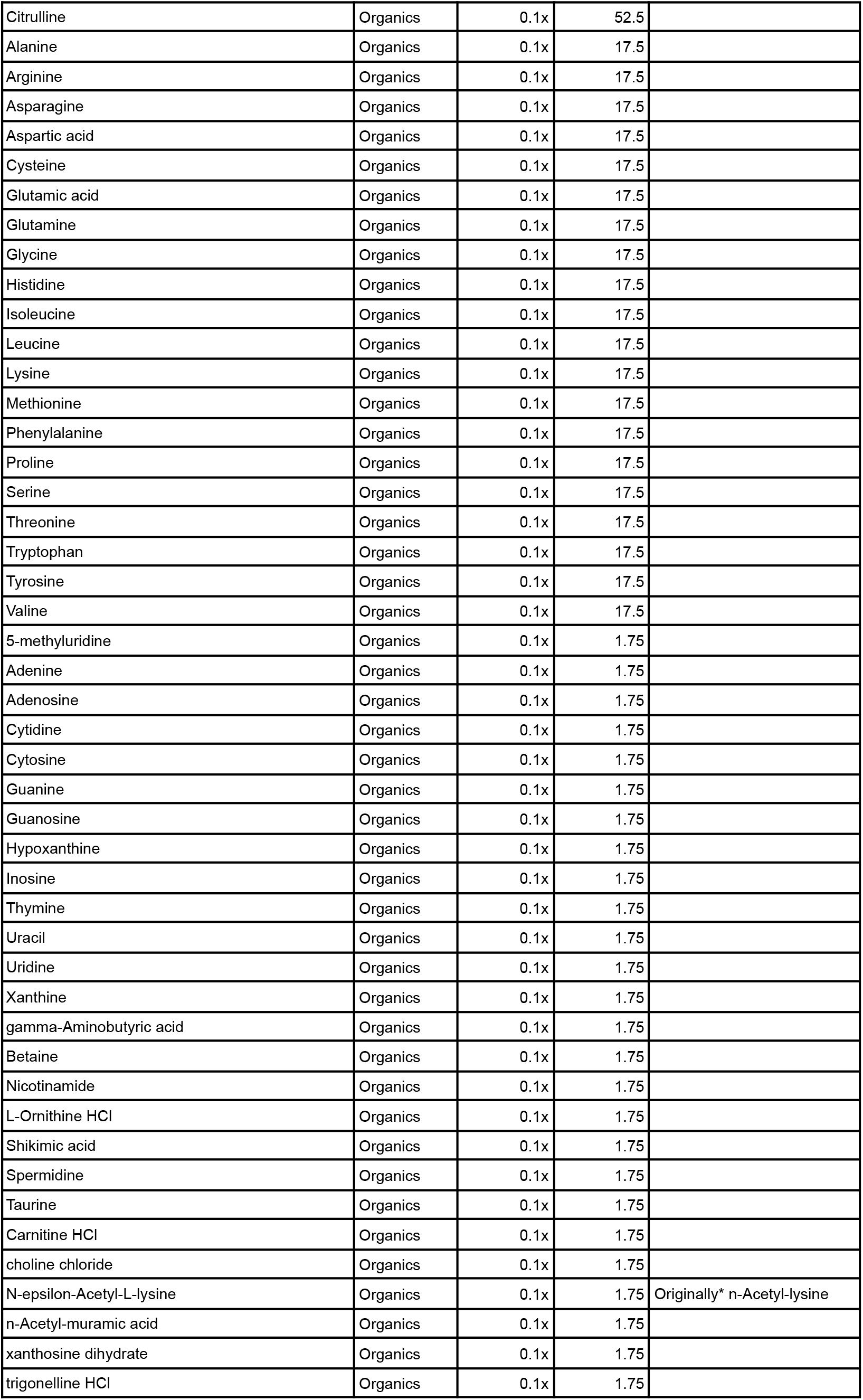
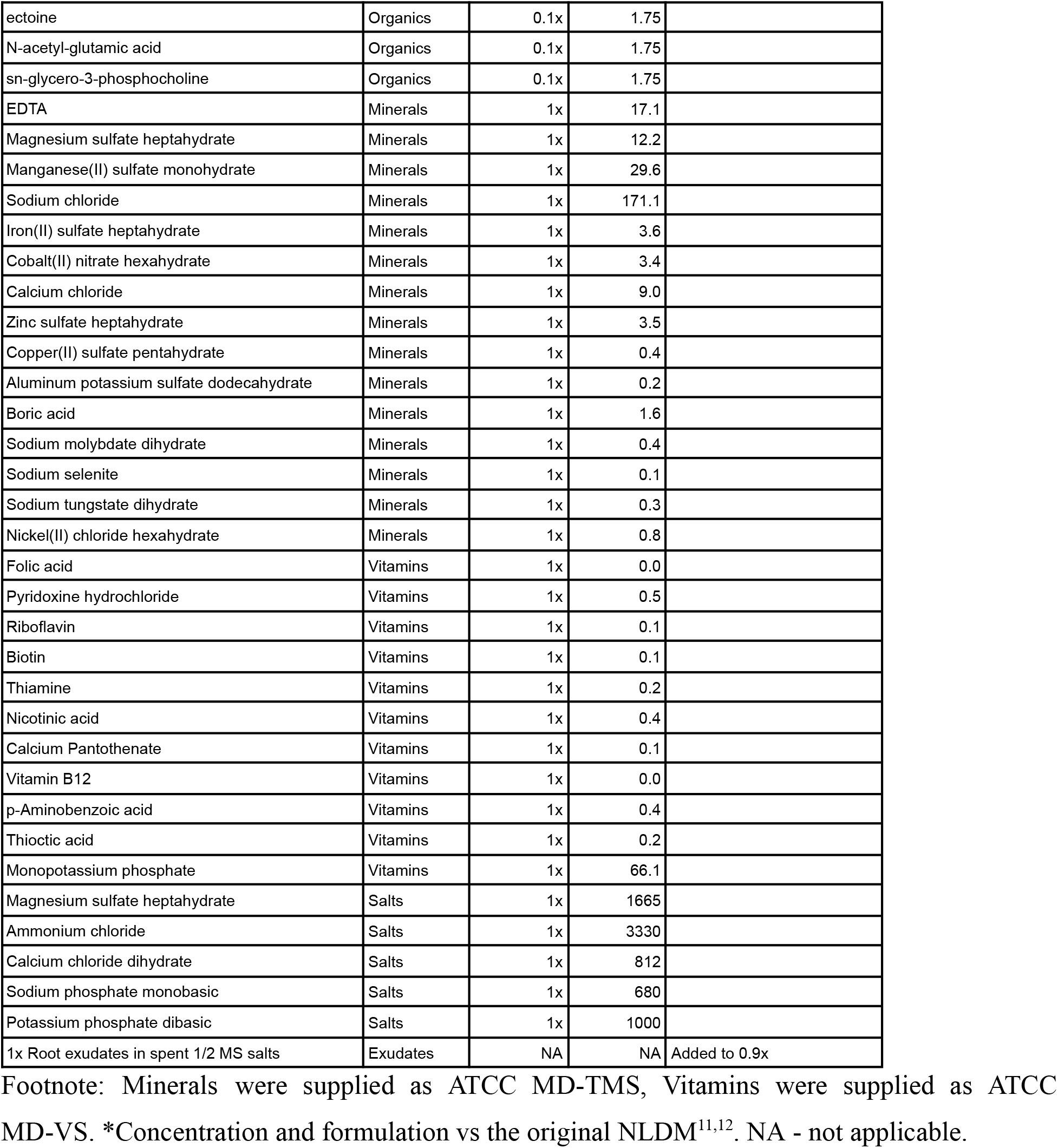
Composition of the exudate-NLDM assay medium.

| Component | Category | Metabolite dilution vs 1xNLDM* | Target metabolite c (uM) | note |
| --- | --- | --- | --- | --- |
| Sucrose | Organics | 0.1x | 87.5 |  |
| D-Glucose | Organics | 0.1x | 87.5 |  |
| D-(+)-Trehalose dihydrate | Organics | 0.1x | 87.5 | Originally* D-(+)-Trehalose |
| Myo-inositol | Organics | 0.1x | 87.5 |  |
| Xylose | Organics | 0.1x | 87.5 |  |
| Mannitol | Organics | 0.1x | 87.5 |  |
| n-Acetyl glucosamine | Organics | 0.1x | 87.5 |  |
| sodium-DL-lactate | Organics | 0.1x | 52.5 | Originally* Sodium-L-Lactate |
| Malic acid | Organics | 0.1x | 52.5 |  |
| Citric acid | Organics | 0.1x | 52.5 |  |
| Succinic acid | Organics | 0.1x | 52.5 |  |
| Sodium Pyruvate | Organics | 0.1x | 52.5 |  |
| alpha Ketoglutaric acid | Organics | 0.1x | 52.5 |  |
| Citrulline | Organics | 0.1x | 52.5 |  |
| Alanine | Organics | 0.1x | 17.5 |  |
| Arginine | Organics | 0.1x | 17.5 |  |
| Asparagine | Organics | 0.1x | 17.5 |  |
| Aspartic acid | Organics | 0.1x | 17.5 |  |
| Cysteine | Organics | 0.1x | 17.5 |  |
| Glutamic acid | Organics | 0.1x | 17.5 |  |
| Glutamine | Organics | 0.1x | 17.5 |  |
| Glycine | Organics | 0.1x | 17.5 |  |
| Histidine | Organics | 0.1x | 17.5 |  |
| Isoleucine | Organics | 0.1x | 17.5 |  |
| Leucine | Organics | 0.1x | 17.5 |  |
| Lysine | Organics | 0.1x | 17.5 |  |
| Methionine | Organics | 0.1x | 17.5 |  |
| Phenylalanine | Organics | 0.1x | 17.5 |  |
| Proline | Organics | 0.1x | 17.5 |  |
| Serine | Organics | 0.1x | 17.5 |  |
| Threonine | Organics | 0.1x | 17.5 |  |
| Tryptophan | Organics | 0.1x | 17.5 |  |
| Tyrosine | Organics | 0.1x | 17.5 |  |
| Valine | Organics | 0.1x | 17.5 |  |
| 5-methyluridine | Organics | 0.1x | 1.75 |  |
| Adenine | Organics | 0.1x | 1.75 |  |
| Adenosine | Organics | 0.1x | 1.75 |  |
| Cytidine | Organics | 0.1x | 1.75 |  |
| Cytosine | Organics | 0.1x | 1.75 |  |
| Guanine | Organics | 0.1x | 1.75 |  |
| Guanosine | Organics | 0.1x | 1.75 |  |
| Hypoxanthine | Organics | 0.1x | 1.75 |  |
| Inosine | Organics | 0.1x | 1.75 |  |
| Thymine | Organics | 0.1x | 1.75 |  |
| Uracil | Organics | 0.1x | 1.75 |  |
| Uridine | Organics | 0.1x | 1.75 |  |
| Xanthine | Organics | 0.1x | 1.75 |  |
| gamma-Aminobutyric acid | Organics | 0.1x | 1.75 |  |
| Betaine | Organics | 0.1x | 1.75 |  |
| Nicotinamide | Organics | 0.1x | 1.75 |  |
| L-Ornithine HCl | Organics | 0.1x | 1.75 |  |
| Shikimic acid | Organics | 0.1x | 1.75 |  |
| Spermidine | Organics | 0.1x | 1.75 |  |
| Taurine | Organics | 0.1x | 1.75 |  |
| Carnitine HCl | Organics | 0.1x | 1.75 |  |
| choline chloride | Organics | 0.1x | 1.75 |  |
| N-epsilon-Acetyl-L-lysine | Organics | 0.1x | 1.75 | Originally* n-Acetyl-lysine |
| n-Acetyl-muramic acid | Organics | 0.1x | 1.75 |  |
| xanthosine dihydrate | Organics | 0.1x | 1.75 |  |
| trigonelline HCl | Organics | 0.1x | 1.75 |  |
| ectoine | Organics | 0.1x | 1.75 |  |
| N-acetyl-glutamic acid | Organics | 0.1x | 1.75 |  |
| sn-glycero-3-phosphocholine | Organics | 0.1x | 1.75 |  |
| EDTA | Minerals | 1x | 17.1 |  |
| Magnesium sulfate heptahydrate | Minerals | 1x | 12.2 |  |
| Manganese(II) sulfate monohydrate | Minerals | 1x | 29.6 |  |
| Sodium chloride | Minerals | 1x | 171.1 |  |
| Iron(II) sulfate heptahydrate | Minerals | 1x | 3.6 |  |
| Cobalt(II) nitrate hexahydrate | Minerals | 1x | 3.4 |  |
| Calcium chloride | Minerals | 1x | 9.0 |  |
| Zinc sulfate heptahydrate | Minerals | 1x | 3.5 |  |
| Copper(II) sulfate pentahydrate | Minerals | 1x | 0.4 |  |
| Aluminum potassium sulfate dodecahydrate | Minerals | 1x | 0.2 |  |
| Boric acid | Minerals | 1x | 1.6 |  |
| Sodium molybdate dihydrate | Minerals | 1x | 0.4 |  |
| Sodium selenite | Minerals | 1x | 0.1 |  |
| Sodium tungstate dihydrate | Minerals | 1x | 0.3 |  |
| Nickel(II) chloride hexahydrate | Minerals | 1x | 0.8 |  |
| Folic acid | Vitamins | 1x | 0.0 |  |
| Pyridoxine hydrochloride | Vitamins | 1x | 0.5 |  |
| Riboflavin | Vitamins | 1x | 0.1 |  |
| Biotin | Vitamins | 1x | 0.1 |  |
| Thiamine | Vitamins | 1x | 0.2 |  |
| Nicotinic acid | Vitamins | 1x | 0.4 |  |
| Calcium Pantothenate | Vitamins | 1x | 0.1 |  |
| Vitamin B12 | Vitamins | 1x | 0.0 |  |
| p-Aminobenzoic acid | Vitamins | 1x | 0.4 |  |
| Thioctic acid | Vitamins | 1x | 0.2 |  |
| Monopotassium phosphate | Vitamins | 1x | 66.1 |  |
| Magnesium sulfate heptahydrate | Salts | 1x | 1665 |  |
| Ammonium chloride | Salts | 1x | 3330 |  |
| Calcium chloride dihydrate | Salts | 1x | 812 |  |
| Sodium phosphate monobasic | Salts | 1x | 680 |  |
| Potassium phosphate dibasic | Salts | 1x | 1000 |  |
| 1x Root exudates in spent 1/2 MS salts | Exudates | NA | NA | Added to 0.9x |

Footnote: Minerals were supplied as ATCC MD-TMS, Vitamins were supplied as ATCC MD-VS. \*Concentration and formulation vs the original NLDM<sup>11,12</sup>. NA - not applicable.

### 2.3. Soil origin and inoculum generation

Soil was collected from Schulenberg Prairie (GPS: 41.81496323898044, −88.09283723716084) in The Morton Arboretum (Lisle, Illinois, USA) in July 2025. We sampled ~1kg of topsoil into a gallon Ziploc Zipper bag from 5 different locations to reflect the variation in moisture and plant communities. After collection, the soil was stored at 4ºC until it was shipped to Lawrence Berkeley National Laboratory. It was shipped with ice packs with overnight shipping. After receiving the soil, it was immediately sieved through a 2 mm stainless steel sieve, stored at 4ºC in the dark in a gallon Ziploc Zipper bag to maintain moisture until inoculum preparation in November 2025. The soil bag was opened regularly during storage to ensure air exchange.

Soil microbial inoculum was prepared under sterile conditions by mixing 1.5 g of soil with 7.5 mL of filter-sterilized Milli-Q water. The soil was vortexed until resuspended and incubated at room temperature for 1 hour. The supernatant was collected by centrifugation at 4°C for 10 minutes at 2500g and used immediately as inoculum.

### 2.4. Microbial culture and sample collection

We mixed 2145 μL of exudate assay media with 55 μL of inoculum (40× inoculum dilution). Uninoculated controls were prepared by replacing the inoculum with 55 μL of sterile Milli-Q water. Each treatment was prepared in three biological replicates. Optical density (OD_600_) was measured (**Fig. 2**) at the start of the culture (t0) by transferring 200 μL aliquots to 96-well plates and measuring absorbance at 600 nm using a Synergy H1 microplate reader (BioTek). The remaining 2000 μL of each mixture were then cultured in autoclaved Corning 24-deep-well plates (VWR 89080-532) sealed with a Breathe-Easy membrane (Sigma-Aldrich, Z380059) and incubated at 650 rpm and 28°C on an Infors HT Multitron Pro. After 5 days of incubation (t+5 d), the final OD_600_ was measured using the same procedure. The remaining culture was then used for downstream metabolomics and microbiome analyses. Specifically, 1 mL of culture was transferred to DNA LoBind tubes (Eppendorf #022431021) and centrifuged at 8000g for 10 min at 4°C. Supernatants (0.85 mL) were collected into new tubes (Eppendorf #022363204) for LC-MS/MS metabolomics, and pellets with residual media (0.15 mL) were retained in DNA LoBind tubes for 16S rRNA gene sequencing. All samples were stored at −80°C prior to analysis.

**Fig. 2:**
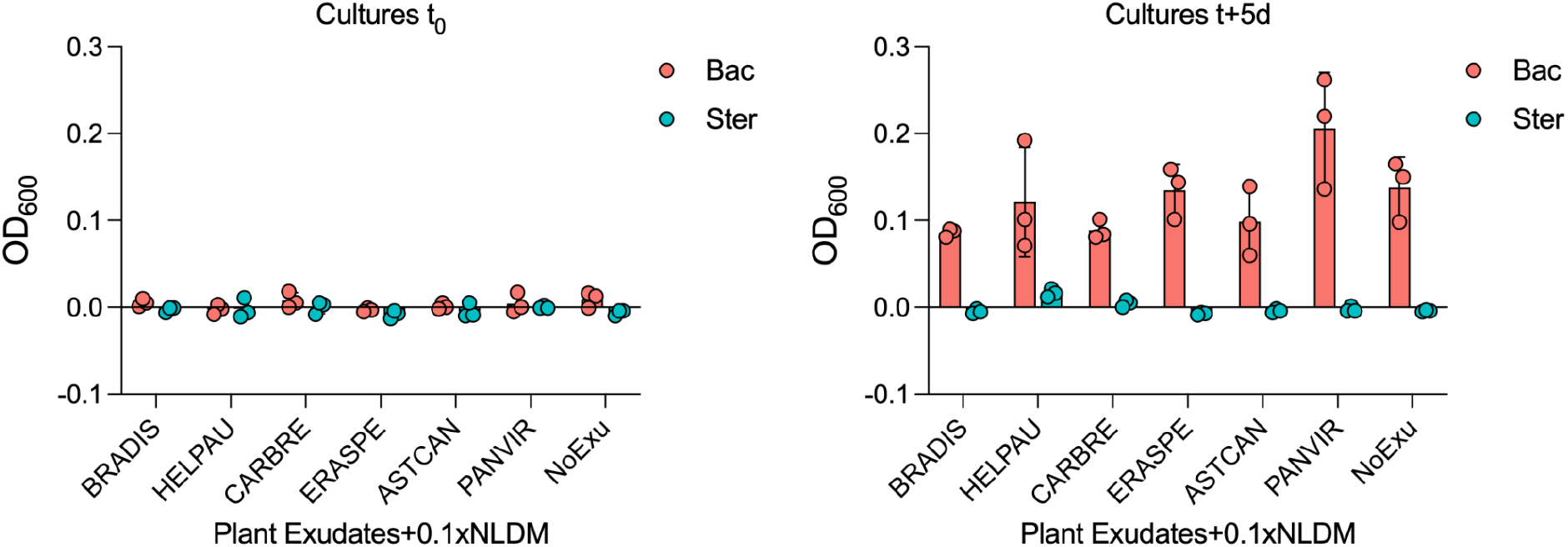
Microbial growth in the culture. Microbial growth in assay media was measured by OD_600_ at t_0_ and t+5 days. The assay media consisted of root exudates mixed with NLDM at 0.1x organics concentration. The bar plot shows the Average ± Standard deviation of the inoculated (Bac) vs. sterile (Ster) treatments; each datapoint is shown (*n*=3). The values were blank subtracted (*n*=54).

### 2.5. LC-MS/MS metabolomics

We prepared soil and spent culture supernatant samples for LC-MS/MS. First, soil was extracted. During extraction, samples were kept on ice, and solvents were chilled at −20 °C. Five g of wet-weight soil were added to 50 mL tubes (525-1109, VWR), including five empty extraction controls. All tubes received 25 mL of LC-MS grade water (9831-03, J.T. Baker), vortexed to ensure the soil was fully saturated and resembled a watery slurry, then sonicated (97043-944, VWR) in ice water for 15 min. Samples were loaded onto a platform shaker (Model SI-1700, Scientific Industries Orbital Genie) to shake at a 45º angle at 200 rpm overnight in a 4 °C cold room (~15 hours). The following morning, samples were centrifuged (5430R, Eppendorf) at 8,000g for 10min at 10 °C, and the supernatant was transferred to a new 50mL tube (525-1109, VWR). Samples were stored at −20 °C for 2 hours, then transferred to the −80 °C freezer to fully freeze.

Frozen spent culture supernatants and soil extracts were lyophilized (7670520, Labconco) for 3 days until completely dry. The dried soil extracts were suspended in 1mL methanol (MX0486, Sigma), vortexed, transferred to a new microcentrifuge tube (022363352, Eppendorf), and sonicated in an ice bath for 15 minutes. Samples were then centrifuged (5430R, Eppendorf) at 10,000g for 5 minutes at 10 °C, and then the supernatant was transferred to a new microcentrifuge tube (022363352, Eppendorf). Supernatants were dried by vacuum concentration (SpeedVac Concentrator, Thermo) and then stored at −80 °C overnight. The following morning, the dried spent culture supernatants and soil extracts were resuspended in 150 µL methanol containing the internal standard mix^19,20^, vortexed, and centrifuged (5430R, Eppendorf) at 10,000g for 5 min at 10 °C. Supernatants were filtered with 0.22 µm PVDF filters (UFC30GVNB, Millipore), transferred to amber glass vials with inserts (5188−6592, Agilent), and sealed with screw caps (5185−5820, Agilent).

The metabolomics samples were analyzed using an LC-MS/MS system consisting of an Agilent 1290 Infinity UHPLC and a Thermo Q Exactive Orbitrap, and analyzed according to established methods for non-polar (C18)^20^ and polar (HILIC)^19^ metabolites in positive and negative modes.

For downstream analysis and archiving, we performed feature-based molecular networking (FBMN)^21^ using GNPS2 (http://gnps2.org)^14^. First, the MZmine 3.7.2^22^ was used to generate a feature list from mzML files utilizing a custom batch workflow. For each feature, the most intense fragmentation spectrum was submitted to GNPS2 for searching against spectral libraries to generate putative annotations. To make overview heatmap figures (**Fig. 3**), the peak area data were downloaded as a feature table from the GNPS2 Analysis Status Page by clicking “Download Quantification file”.

**Fig. 3:**
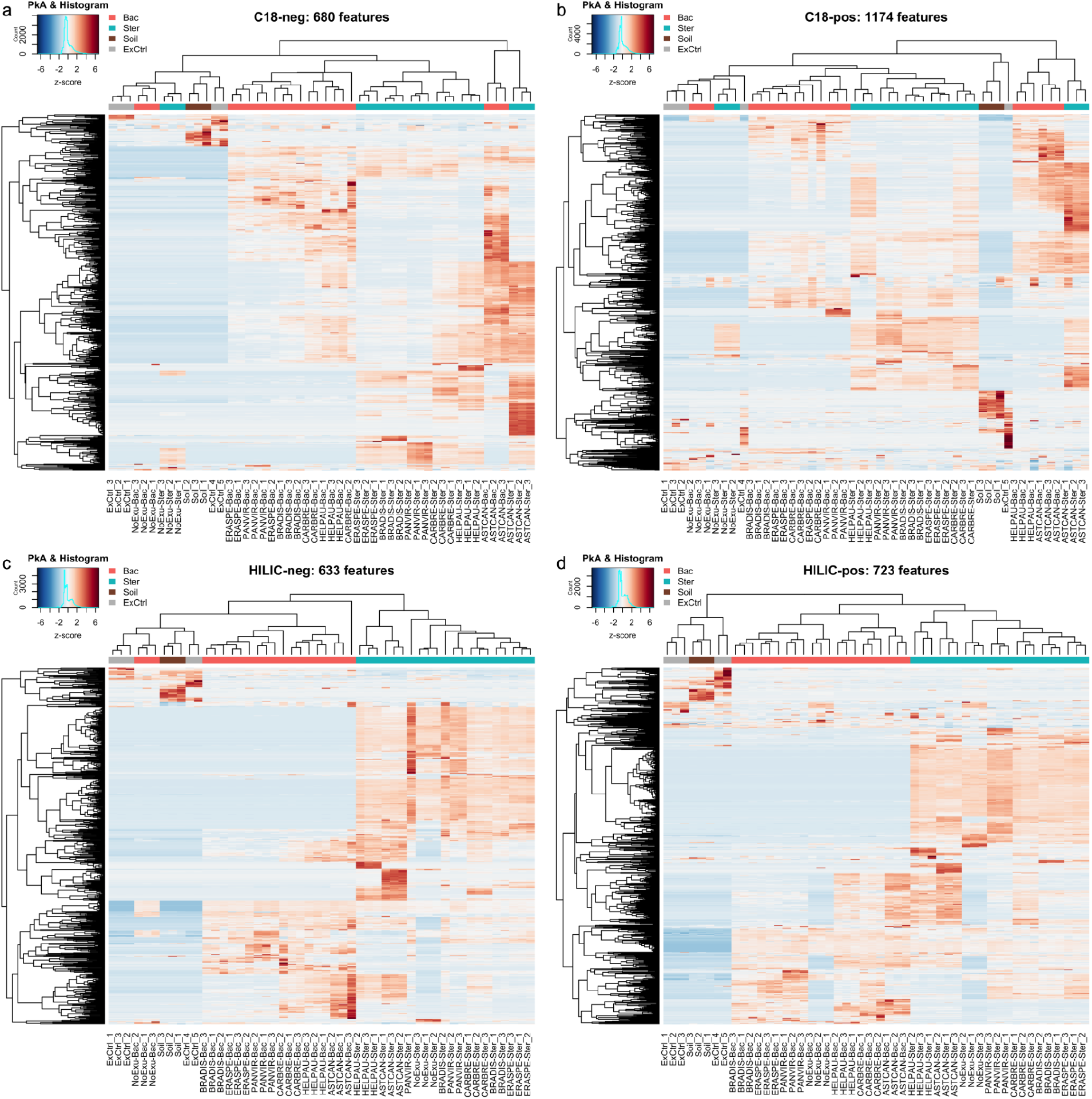
Metabolomics data overview shown as a heatmap of all samples for detected features in GNPS2. Peak areas of features are shown as z-scores. The raw peak area data were downloaded from the GNPS2 Analysis page as a quantification file. Each panel represents a different acquisition mode (**a**) C18-neg, (**b**) C18-pos, (**c**) HILIC-neg, (**d**) HILIC-pos. The number of rows in each heatmap represents the number of features.

### 2.6. Microbiome amplicon sequencing and data analysis

The pellets from the inoculated (Bac) treatment were processed and analyzed with the Microbiome Sequencing Service: Full-length 16S Amplicon Sequencing (Zymo Research, Irvine, CA). The DNA was extracted with ZymoBIOMICS®-96 MagBead DNA Kit (Zymo Research, Irvine, CA). Full-length 16S sequencing libraries were prepared according to the Quick-16STM Full-Length Library Prep Kit (Zymo Research, Irvine, CA). In brief, the whole 16S gene was amplified using the 27f (AGRGTTYGATYMTGGCTCAG) and 1492r (RGYTACCTTGTTACGACTT) primers with barcodes and adapters. Two nanograms of DNA were used as the PCR input for each sample. The PCR was run using the kit-provided premix for 35 cycles on the iconPCR® instrument (n6Tec, Pleasanton, CA) according to the protocol. The subsequent normalized PCR products from each reaction were pooled and cleaned up with Select-a-Size DNA Clean & Concentrator− (Zymo Research, Irvine, CA) to retain fragments greater than 300 bp. The final pooled library was quantified with TapeStation® (Agilent Technologies, Santa Clara, CA) and Qubit® (Thermo Fisher Scientific, Waltham, WA). To prepare for PacBio® Sequencing, the pooled library was further processed using the Kinnex 16S rRNA Kit® (Pacific Biosciences, Menlo Park, CA).

The ZymoBIOMICS® Microbial Community Standard (Zymo Research, Irvine, CA) was used as a positive control for each DNA extraction. The ZymoBIOMICS® Microbial Community DNA Standard (Zymo Research, Irvine, CA) was used as a positive control for each targeted library preparation. Negative controls (i.e., blank extraction control and blank library preparation control) were included to assess the level of bioburden introduced by the wet-lab process. The library was sequenced on a single 8M SMRT cell using the PacBio Sequel IIe system for 15 hours. HiFi reads, produced by Circular Consensus Sequencing mode, were generated by the sequencer.

Unique amplicon sequence variants were inferred from raw reads using the DADA2 pipeline ^23^. Potential sequencing errors and chimeric sequences were also removed with the Dada2 pipeline. Taxonomy assignment was performed using UCLUST from Qiime v.1.9.1 with the Zymo Research Database, an internally designed and curated 16S database, as a reference. Composition visualization, alpha-diversity, and beta-diversity analyses were performed with Qiime v.1.9.1^24^. If applicable, taxonomies with significant abundance across groups were identified by LEfSe^25^ using default settings. Other analyses, such as heatmaps, Taxa2ASV Deomposer, and PCoA plots, were performed with internal scripts by Zymo Research.

### 2.7. Software

Heatmap plots were generated in R (version 4.5.2) using the heatmap.2 function from the gplots package^26^. Biorender.com was used to create graphical overviews. Microsoft Excel (version 16.108.1) was used to store and manipulate data frames.

## 3. Data Records

The datasets generated in this study are distributed across multiple repositories, including NCBI, Figshare, MassIVE, and GNPS2 (**Fig. 1**). The raw files from full-length 16S rRNA sequencing were deposited in FASTQ format to the Sequence Read Archive (SRA) at NCBI (https://www.ncbi.nlm.nih.gov) under BioProject PRJNA1460902.

Figshare serves as the primary repository for sample information, R code, metabolomics processing files, and sequencing outputs, available as a collection at https://doi.org/10.6084/m9.figshare.c.8447566. The metadata overview table (.xlsx) provides sample identifiers, OD_600_ measurements, and naming conventions that link samples across metabolomics and microbiome datasets, along with sample submission sheets for the respective omics analyses. A separate folder, “16S-seq_Zymo,” contains microbiome composition data from full-length 16S rRNA sequencing conducted by Zymo Research. This folder includes a zipped report with bioinformatic analyses, including compositional bar plots, heatmaps, alpha- and beta-diversity metrics, and LEfSe statistical group comparisons. Two accompanying PDF documents are provided: (i) a description of the sequencing service and (ii) a folder navigation guide to facilitate interpretation of the report structure and outputs. Additionally, a sequencing sample submission sheet (.xlsx) is included. The Figshare collection also includes MZmine batch parameter files (.xml) in the folder “MZmine-batch-params” for each LC-MS/MS acquisition mode, enabling the re-creation of the feature list for downstream analysis in GNPS2. The “HeatmapR” folder contains R scripts for generating metabolomics heatmaps, along with accompanying spreadsheets (.xlsx) containing peak area data for each acquisition mode used as input.

The LC-MS files in .raw format are stored at MassIVE (https://massive.ucsd.edu/) under ID number MSV000101683 and can be accessed via https://doi.org/10.25345/C5T727W15. GNPS2 is used to store LC-MS/MS files in .mzML format and for downstream bioinformatic analysis. There is extensive documentation for the GNPS2 platform at https://gnps2.org/. We have utilized Feature-Based Molecular Networking^21^ that allows for visualization of the data at the GNPS2 Analysis Status Page for each of the acquisition modes: C18-neg: https://gnps2.org/status?task=b99cf9be1b1c44b885470f620bb78820, C18-pos: https://gnps2.org/status?task=543fd0e3aa8c4338803d5ac7467d697d, HILIC-neg: https://gnps2.org/status?task=068d14aeb8c24ba0a12fe78644874404, HILIC-pos: https://gnps2.org/status?task=0bd6052f85044e58bc24ca507e644eb8.

## 4. Technical validation

The OD_600_ measurement was used to confirm microbial growth in inoculated cultures relative to sterile media, ensuring the samples were fit for downstream analyses (**Fig. 2**).

Metabolomics used checks at multiple levels to ensure technical accuracy in the experimental design and LC-MS/MS analysis. First, we included extraction and technical controls in our experimental design. Technical controls (TxCtrl) consisted of incubated sterile (uninoculated) media for each treatment group, serving as a baseline. Extraction controls (ExCtrl) are empty tubes processed in the same way as the TxCtrl and sample tubes to control for contaminations. The technical validations for LC-MS/MS analysis are detailed in the protocols ^19,20^. Briefly, quality control (QC) injections containing a defined mixture of compounds with known m/z, retention times, and MS/MS spectra were included to monitor instrument performance, including mass accuracy, signal intensity, and retention time stability. Methanol blanks were used to assess background signal and potential carryover. In addition, a mixture of isotopically labeled internal standards was added to all samples and injected independently throughout the run to monitor instrument performance and assess sample-level variability, including retention time shifts, injection consistency, matrix effects, and approximate quantitation of unlabeled molecules.

Microbiome data quality was supported through standardized DNA extraction, library preparation, and sequencing workflows with appropriate controls. Positive controls, including defined microbial community and DNA standards, were included at the extraction and library preparation stages to verify performance and taxonomic accuracy, while negative controls (blank extraction and library controls) assessed background contamination. Sequencing generated high-fidelity reads, and downstream processing included denoising, chimera removal, and inference of amplicon sequence variants using established pipelines to remove sequencing errors and artifacts prior to taxonomic assignment.

## 5. Data availability

The FASTQ sequencing files were uploaded to the SRA at NCBI (https://www.ncbi.nlm.nih.gov) under PRJNA1460902. The 16S rRNA sequencing report by Zymo was uploaded into Fishare as a collection at https://doi.org/10.6084/m9.figshare.c.8447566, alongside metadata, R code, and MZmine batch files. The metabolomics files in .raw format are deposited in MassIVE (https://massive.ucsd.edu/) under ID number MSV000101683 and can be accessed via https://doi.org/10.25345/C5T727W15. The untargeted metabolomics can be accessed via GNPS2 Analysis Status Page: C18-neg: https://gnps2.org/status?task=b99cf9be1b1c44b885470f620bb78820, C18-pos: https://gnps2.org/status?task=543fd0e3aa8c4338803d5ac7467d697d, HILIC-neg: https://gnps2.org/status?task=068d14aeb8c24ba0a12fe78644874404, HILIC-pos: https://gnps2.org/status?task=0bd6052f85044e58bc24ca507e644eb8

## 6. Code Availability

The R code and data used to generate the metabolomics heatmaps are deposited in Figshare at https://doi.org/10.6084/m9.figshare.c.8447566.

## 8. Author Contributions

VN: Conceptualization, Methodology, Investigation, Data Curation, Visualization, Writing – Original Draft. MdR: Methodology. AV: Methodology. VS: Investigation. MM: Resources. CH: Investigation. NAB: Methodology, Funding Acquisition. TRN: Supervision, Funding Acquisition. All authors: Writing – Review & Editing.

## 10. Acknowledgments

We acknowledge Felicia Wong for technical assistance with experiments. We would like to thank Thomas Harwood for technical guidance regarding the GNPS2 workflow. Data analysis utilized resources from the National Energy Research Scientific Computing Center (NERSC), a DOE Office of Science User Facility (Contract No. DE-AC02-05CH11231)

## References

1. Zhalnina, K. et al.. Dynamic root exudate chemistry and microbial substrate preferences drive patterns in rhizosphere microbial community assembly. Nat. Microbiol. 3, 470–480 (2018).

2. Sasse, J., Martinoia, E. & Northen, T. Feed your friends: do plant exudates shape the root microbiome? Trends Plant Sci. 23, 25–41 (2018).

3. McLaughlin, S., Zhalnina, K., Kosina, S., Northen, T. R. & Sasse, J. The core metabolome and root exudation dynamics of three phylogenetically distinct plant species. Nat. Commun. 14, 1649 (2023).

4. Williams, A. et al.. Root functional traits explain root exudation rate and composition across a range of grassland species. J. Ecol. (2021) doi:10.1111/1365-2745.13630.

5. Ding, Y. et al.. Insights into the role of dopamine in rhizosphere microbiome assembly. bioRxiv (2024).

6. Novak, V. et al.. Reproducible growth of Brachypodium in EcoFAB 2.0 reveals that nitrogen form and starvation modulate root exudation. Sci. Adv. 10, eadg7888 (2024).

7. Bai, Y. & Cotrufo, M. F. Grassland soil carbon sequestration: Current understanding, challenges, and solutions. Science 377, 603–608 (2022).

8. Ulbrich, T. C., Rivas-Ubach, A., Tiemann, L. K., Friesen, M. L. & Evans, S. E. Plant root exudates and rhizosphere bacterial communities shift with neighbor context. Soil Biol. Biochem 172, 108753 (2022).

9. Oburger, E. & Jones, D. L. Sampling root exudates – Mission impossible? Rhizosphere 6, 116–133 (2018).

10. Novak, V. et al.. Breaking the reproducibility barrier with standardized protocols for plant-microbiome research. PLoS Biol. 23, e3003358 (2025).

11. de Raad, M. et al.. A defined medium for cultivation and exometabolite profiling of soil bacteria. Front. Microbiol. 13, 855331 (2022).

12. De Raad, M. NLDM: A defined medium for cultivation and exometabolite profiling of soil bacteria v1. (2022) doi:10.17504/protocols.io.14egn7o2yv5d/v1.

13. Jenkins, S. et al.. Construction of viable soil defined media using quantitative metabolomics analysis of soil metabolites. Front. Microbiol. 8, 2618 (2017).

14. Wang, M. et al.. Sharing and community curation of mass spectrometry data with Global Natural Products Social Molecular Networking. Nat. Biotechnol. 34, 828–837 (2016).

15. Sayers, E. W. et al.. Database resources of the National Center for Biotechnology Information in 2025. Nucleic Acids Res. 53, D20–D29 (2025).

16. Wilkinson, M. D. et al.. The FAIR Guiding Principles for scientific data management and stewardship. Sci. Data 3, 160018 (2016).

17. Kelliher, J. M. et al.. STREAMS guidelines: standards for technical reporting in environmental and host-associated microbiome studies. Nat. Microbiol. 10, 3059–3068 (2025).

18. Schoch et al. NCBI Taxonomy: a comprehensive update on curation, resources and tools. Database 2020, (2020).

19. B. Louie, K. et al. JGI/LBNL Metabolomics - Standard LC-MS/MS ESI Method - Polar HILIC-Z v1. (2024) doi:10.17504/protocols.io.kxygxydwkl8j/v1.

20. B. Louie, K. et al. JGI/LBNL Metabolomics - Standard LC-MS/MS ESI Method – Nonpolar C18 v1. (2024) doi:10.17504/protocols.io.261ge5mzjg47/v1.

21. Nothias, L.-F. et al.. Feature-based molecular networking in the GNPS analysis environment. Nat. Methods 17, 905–908 (2020).

22. Schmid, R. et al.. Integrative analysis of multimodal mass spectrometry data in MZmine 3. Nat. Biotechnol. 41, 447–449 (2023).

23. Callahan, B. J. et al.. DADA2: High-resolution sample inference from Illumina amplicon data. Nat. Methods 13, 581–583 (2016).

24. Caporaso, J. G. et al.. QIIME allows analysis of high-throughput community sequencing data. Nat. Methods 7, 335–336 (2010).

25. Segata, N. et al.. Metagenomic biomarker discovery and explanation. Genome Biol. 12, R60 (2011).

26. Warnes, G. R. et al.. gplots: Various R Programming Tools for Plotting Data. R package version 3.1.3.1. https://CRAN.R-project.org/package=gplots (2024).

